# Disentangling causal relationships between inflammatory markers and depression: a bidirectional Mendelian randomization analysis

**DOI:** 10.1101/712133

**Authors:** Christina Dardani, James Yarmolinsky, Jamie Robinson, Jie Zheng, George Davey Smith, Sarah J Lewis, Lindsey I Sinclair

**Author notes:** Joint first authors (Christina Dardani, James Yarmolinsky). **Corresponding author address**: Dr Lindsey Sinclair, Dementia Research Group, Level 1 Learning & Research Building, Southmead Hospital, Bristol, BS_10 5_NB • (0117) 4147807 •.

## Abstract

**Background:** The inflammatory markers C-reactive protein (CRP), interleukin-1 receptor antagonist (IL1-Ra), and interleukin-6 (IL-6) have been associated with depression risk in observational studies. The causal nature of these associations is unclear as conventional observational designs are susceptible to reverse causation and residual confounding. Bidirectional Mendelian randomization (MR) analysis uses genetic variants to proxy for risk factors to help elucidate the presence, magnitude, and direction of causal relationships between traits.

**Methods:** We performed bidirectional two-sample MR to examine causal associations between circulating CRP, IL1-Ra, and IL-6 and major depressive disorder (MDD) in 135,458 cases and 344,901 controls in the Psychiatric Genetics Consortium. Genetic instruments to proxy inflammatory markers and liability to MDD were constructed by obtaining single-nucleotide polymorphisms (SNPs) associated with these phenotypes in genome-wide association study meta-analyses. Wald ratios and inverse-variance weighted random-effects models were employed to generate causal effect estimates and various sensitivity analyses were performed to examine violations of MR assumptions.

**Results:** There was evidence supporting a causal effect of circulating IL-6 on risk of MDD (per natural-log increase: OR 0.85, 95% CI: 0.75-0.96, *P*=0.007). Higher circulating levels of IL-6 as influenced by variants in the IL6R gene region represent lower cellular binding of IL-6 to its receptor and therefore the present results suggest that IL-6 increases the risk of MDD. We found limited evidence supporting a causal effect of CRP (1.06, 95% CI 0.93-1.22; P=0.36) or IL1-Ra (OR 0.95, 95% CI: 0.87-1.03, P=0.20) on risk of MDD. Reverse direction MR analyses suggested limited evidence for a causal effect of genetic liability to MDD on any of the inflammatory markers examined.

**Conclusions:** These findings support a causal role of IL-6-related pathways in development of major depressive disorder and suggest the possible efficacy of interleukin-6 inhibition as a therapeutic target for depression.

## INTRODUCTION

Depression is a common mental disorder with an estimated lifetime prevalence of 13-18%^1,2^. Despite major advances in the field, current antidepressant treatment options are still suboptimal; around one third of patients appear to be resistant to treatment^3^, and virtually all antidepressant drugs have a delayed onset of action^4^. This has a severe impact on the affected individuals’ quality of life and it imposes large societal costs^5,6^. Enhancing current understanding of the aetiopathogenic factors implicated in this complex condition is an important step towards offering new treatment options.

One promising area of research into the aetiology of depression is the inflammatory response. In the 1990s it was first proposed that neuroinflammation may be a cause of depression^7^. Since then, additional evidence to suggest a link between the condition and inflammation has continued to emerge. Observational studies indicate an increased prevalence of depressive disorders in individuals with chronic inflammatory conditions^8,9^. Further, in a recent meta-analysis of 82 case-control studies, elevated levels of serum circulating cytokines, among them: interleukin-1 receptor antagonist (IL-1Ra) and interleukin-6 (IL-6), were found in individuals diagnosed with depression as compared to healthy controls^10^. Additionally, a small number of prospective studies have reported associations between elevated serum CRP, IL-1Ra and IL-6 and the later development of depression in both young adult and geriatric populations^11–14^.

However, there are important challenges when appraising the potential causal role of inflammatory markers in depression from observational studies. This is because conventional epidemiological study designs, including case-control studies and prospective studies with insufficient follow-up time, may fail to distinguish whether the inflammatory response is a cause or consequence of depression (“reverse causation”)^15,16^. Further, such analyses are prone to residual confounding due to unmeasured or imprecisely measured confounders (i.e., common causes of inflammation and depression that are not on the causal pathway between these traits).

One method to overcome the limitations imposed on causal inference from conventional observational studies is Mendelian randomization (MR). MR is an instrumental variables (IV) approach which utilizes germline genetic variants as proxies for risk factors to assess the causal effects of these factors on disease outcomes^17,18^. On this basis, the method is effective in minimising confounding bias and is immune to reverse causation since genetic variants are randomly assorted at meiosis and are fixed at conception^17,19^.

To date, MR studies investigating associations between circulating inflammatory markers and depression have been inconclusive. In a Danish general population sample there was limited evidence to suggest a causal effect of genetically proxied CRP on either hospitalisation with depression (N cases=1,145) or antidepressant medication use (N cases= 8,621)^20^. By contrast, in a subsample of the UK Biobank study participants presenting with probable moderate/severe lifetime depression, as defined by self-report (N cases= 14,701), there was strong evidence of a positive causal effect of both CRP and IL-6 on risk of depression^21^. However, deriving conclusions on the possible causal effects of inflammatory markers on risk of depression when synthesising these studies is challenging given: i. the limited to modest sample size of some of the studies, e.g.^20,22^, ii. variable outcome definitions employed (e.g. depressive episode, depressive disorder^20^, probable moderate or severe lifetime major depression^21^), and iii. differing methods of case ascertainment e.g. self-reports^21^ vs registry-based data^20^.

In this context, we aimed to investigate the presence, magnitude, and direction of putative causal relationships between circulating inflammatory markers and depression through two-sample MR^23^ using the largest genotyped sample of cases diagnosed with major depressive disorder. We investigated the bidirectional causal effects of: circulating C-reactive protein (CRP), Interleukin-1 Receptor antagonist (IL-1Ra) and Interleukin-6 (IL-6) and risk of major depressive disorder in 135,458 cases and 344,901 controls in the Psychiatric Genetics Consortium^24^.

## METHODS

### Two-sample Mendelian randomization

MR can be used to generate unbiased causal estimates of the effects of exposures on disease outcomes under specific assumptions: the IV (here, one or more genetic variants) is: i) robustly associated with the exposure of interest; ii) not associated with any confounding factor(s) that would otherwise distort the association between the exposure and outcome; and iii) not associated with the outcome, independent of changes in the exposure (“exclusion restriction criterion”)^25^. Violation of one or more of these assumptions means that instruments are invalid and, consequently, that findings from such an analysis may yield a biased effect estimate.

In order to overcome limitations of previous MR studies in the field, we applied two-sample MR. In two-sample MR, instrument-exposure and instrument-outcome effect sizes and standard errors are extracted from summary genetic data from two independent samples representative of the same underlying population. Such an approach allows statistical power and precision of causal estimates to be increased because it does not require measured exposure, outcome and genotyped data on all participants within a single sample^19,26^ and can therefore utilize summary effect estimates from previously-published large genome-wide association studies (GWAS).

### GWAS Summary Data: Inflammatory Markers

For analyses examining the effect of inflammatory markers on MDD, summary genetic association data for inflammatory markers were extracted from published analyses as follows: serum CRP data were extracted from a GWAS meta-analysis of 88 studies including 204,402 participants^27^; circulating IL-6 data were obtained from a GWAS meta-analysis of 40 studies including up to 133,449 individuals; and serum IL-1Ra data were obtained from a GWAS of 6,135 individuals^28^. (1) All analyses were adjusted for age, sex, and population sub-structure. For reverse direction MR analyses (i.e., examining the effect of genetic liability to MDD on inflammatory marker levels), complete summary genetic association data were not available for SNPs used to proxy MDD from all of these studies. Consequently, data for plasma CRP, IL-6, and IL-1Ra were obtained from a GWAS meta-analysis of 3,301 individuals of European descent, adjusted for age, sex, duration between blood draw and processing, and principal components of ancestry^29^. Extensive information regarding the GWAS analyses included can be found in the original publications.

### GWAS Summary Data: Major Depressive Disorder

We obtained summary genetic association data from analyses in 135,458 major depressive disorder (MDD) cases and 344,901 controls of European ancestry in the Psychiatric Genetics Consortium (PGC)^24^. The PGC consisted of a combined meta-analysis of 29 individual cohorts forming the current PGC samples (PCG29 cohort: 6,823 MDD cases; 25,632 controls) and six independent cohorts (118,635 cases; 319,269 controls). Cases in the PCG29 cohort were defined using international consensus criteria (DSM-IV, ICD-9, or ICD-10) for a lifetime diagnosis of MDD using structured diagnostic instruments from assessments by trained interviewers, clinician-administered checklists, or medical record review and controls in 22/29 samples were screened for the absence of lifetime MDD and randomly selected from the population. In the remaining cohorts, controls were not screened for MDD. Cases in the additional six independent cohorts used a range of methods for defining MDD. Most studies included applied typical inclusion and exclusion criteria for both cases and controls, for example, the exclusion of individuals with other mental disorders such as bipolar disorder or schizophrenia. Further information on the GWAS can be found in the original publication.

### Genetic instrument construction: Inflammatory Markers

As genetic instruments for inflammatory markers, we used one of two approaches:

i. we selected individual cis-acting variants (located within proximity to the protein-coding gene) robustly (*P* < 5 × 10^−8^) associated with the marker (**IL-1Ra**) or multiple weakly correlated (r^2^ ≤ 0.41) variants located within proximity to *IL6R* that were associated (*P* < 5 × 10^−7^) with the marker (**IL-6**),
ii. we constructed multi-allelic instruments consisting of *cis*- and *trans*-acting independent (r^2^ < 0.001) variants that were associated (*P* < 5 × 10^−8^) with the inflammatory marker (**CRP**).

Characteristics of the genetic instruments used to proxy inflammatory markers are presented in Supplementary Table 1.

### Genetic instrument construction: Major Depressive Disorder

To examine genetic liability to MDD, we selected 36 SNPs shown to robustly (*P* < 5 × 10^−8^) and independently (r^2^ <0.001) associate with MDD within the PGC GWAS meta-analysis.

### Statistical analysis

#### IL-1Ra

The risk factor had one SNP as an instrument. The Wald ratio was used to generate effect estimates and the delta method was used to approximate standard errors^30^.

#### IL-6

The risk factor had three weakly correlated SNPs combined into a multi-allelic instrument. Causal estimates were generated using a modified random-effects inverse-variance weighted (IVW) model that inflates standard errors to account for these correlations^31^. Correlation matrices were obtained from the 1000 Genomes Phase 3 (European) panel^32^.

#### CRP

The risk factor had 30 independent SNPs as instruments. Causal estimates were generated using inverse-variance weighted (IVW) random-effects models^33^.

#### Sensitivity Analyses

For CRP and genetic liability to depression analyses, the following sensitivity analyses were performed to examine evidence of horizontal pleiotropy (a single locus influencing an outcome through one or more biological pathways independent to that of the presumed exposure); a violation of the “exclusion restriction criterion”): MR-Egger regression, weighted median, and weighted mode estimates^33–35^.

MR Egger is similar to the IVW, performing a weighted generalised linear regression of the SNP-outcome coefficients on the SNP-exposure coefficients, but unlike the IVW, has an unconstrained intercept term. This way, under the assumptions: i. the strength of the SNP-exposure associations is independent of the degree of horizontal pleiotropy (InSIDE assumption), and ii. the measurement error in the instrument is negligible (NOME assumption), the intercept term of the MR Egger provides an estimate of the directional pleiotropic effects (unbalanced horizontally pleiotropic effects) of the SNPs on the outcome, with a non-zero intercept indicating directional pleiotropy. The beta coefficient of the MR Egger regression represents an estimate of the causal effect of an exposure on an outcome accounting for directional pleiotropic effects.

The weighted median provides an estimate of the median of the weighted empirical distribution function of Wald ratio estimates and assumes that at least 50% of the information in a multi-allelic instrument stems from SNPs that are valid instruments.

The weighted mode assumes that the most common effect estimate of the instrumental variables stems from valid instruments and generates a causal effect estimate using the weighted mode (weighted by the inverse variance of the SNP-outcome association) of a smoothed empirical density function of the individual instrumental variables.

For CRP analyses, we additionally generated causal effect estimates using an instrument consisting of four weakly-correlated (r^2^ ≤ 0.20) cis-acting variants within *CRP* using a modified random-effects IVW model that accounts for correlations between SNPs.

For IL-6 analyses, we also evaluated the causal effect of soluble IL-6 receptor (sIL-6R) on risk of MDD. A single SNP in *IL-6R* identified from a collaborative meta-analysis of 125,222 participants was employed as instrument^36^. The Wald ratio was used to generate the causal effect estimate of sIL-6R on risk of MDD. Further details on the instrument can be found in Supplementary Table 1.

For inflammatory markers showing evidence of association with MDD (*P* <0.05), we also tested whether the genetic instrument employed to proxy the inflammatory marker was associated with previously identified causal risk factors for depression: i. body mass index^37^, ii. neuroticism^38^, and iii. physical activity^39^. GWAS data on each risk factor were selected on the basis of previous MR studies identifying causal associations with MDD^37–39^. Further information on the specific GWAS studies utilized can be found in the Supplementary Material. When evidence for a causal effect was identified between an inflammatory marker and i. body mass index, ii. neuroticism, iii. physical activity, we additionally performed multivariable MR. The method is an extension of MR, in which multiple exposures are entered within the same model and their independent effects on the outcome can be estimated. Detailed information on the method can be found elsewhere^40^.

Lastly, in cases that an exposure was showing evidence of a causal role (*P* <0.05), we performed iterative leave-one-out permutation analysis to examine whether any of the results were driven by any individual SNP.

All statistical analyses were performed using R version 3.3.

## RESULTS

### The causal effect of inflammatory markers on MDD

In Mendelian randomization analyses examining the effect of circulating inflammatory markers on MDD, circulating interleukin-6 had a causal effect on risk of major depressive disorder (per natural-log increase in interleukin-6 (ng/mL): OR 0.85, 95% CI: 0.75-0.96, *P*=0.007). Results for interleukin-6 were consistent when performing iterative leave-one-out analyses (Supplementary Table 2). Higher circulating levels of IL-6 as proxied by genetic variants in/near *IL6R* represent lower cellular binding of IL-6 to its receptor and therefore these findings support a role of interleukin-6 in risk of MDD. In sensitivity analyses examining the effect of sIL-6R, there was likewise evidence for an effect of this marker on risk of MDD (per natural-log increase in sIL-6R: OR 1.06, 95%CI: 1.01- 1.12; *P*= 0.04).

There was little evidence that C-reactive protein (per natural-log increase in CRP (mg/L): OR 1.06, 95% CI: 0.93-1.22; *P*=0.36), or interleukin-1 receptor antagonist (per natural-log increase (pg/mL): OR 0.95, 95% CI: 0.87-1.03, *P*=0.20) had a causal effect on risk of major depressive disorder. Sensitivity analyses for CRP were consistent when employing MR-Egger regression, a weighted median estimator, a weighted mode estimator, and when using an instrument restricted to genetic variants located within CRP (OR: 0.99, 95% CI: 0.92-1.07; *P*=0.82) (Table 1).

**Table 1.** Causal estimates of circulating inflammatory markers on risk of major depressive disorder in the Psychiatric Genetics Consortium

| Inflammatory marker | IVW/Wald ratio<br>OR (95% CI)<br>*P*-value | MR-Egger<br>OR (95% CI)<br>*P*-value | Weighted median<br>OR (95% CI)<br>*P*-value | Weighted mode<br>OR (95% CI)<br>*P*-value |
| --- | --- | --- | --- | --- |
| CRP (mg/L) | 1.06 (0.93-1.22)<br>0.36 | 1.16 (0.73-1.83)<br>0.54 | 0.99 (0.83-1.17)<br>0.88 | 0.95 (0.72-1.26)<br>0.72 |
| IL1-Ra (pg/mL) | 0.95 (0.87-1.03)<br>0.20 | - | - | - |
| IL-6 (ng/mL) | 0.85 (0.75-0.96)<br>0.007 | - | - | - |

IVW = Inverse-variance weighted, OR = Odds Ratio, 95% CI = 95% Confidence Interval, CRP = C-reactive protein, IL1-Ra = Interleukin-1 Receptor antagonist, IL-6 = Interleukin-6. OR represents the exponential change in the odds of MDD per each unit increase in natural log-transformed inflammatory marker.

### The effect of IL-6 on causal risk factors for MDD

There was little evidence to suggest a causal effect of IL-6 (ng/mL) on BMI (per natural-log increase: b= −0.027, 95% CI: −0.15−0.09; *P*=0.647) or physical activity as measured by mean acceleration (per natural-log increase: b= 0.46, 95% CI: −0.31-1.23; *P*=0.240). However, we found evidence suggesting a causal effect of IL-6 on neuroticism (per natural-log increase: OR 1.07, 95% CI: 1.02-1.12; *P*=0.011) (Supplementary table 3). The result was consistent when performing iterative leave-one-out analyses (Supplementary Table 4). Multivariable MR suggested that the effect of IL-6 on risk of MDD was largely independent of neuroticism (per natural-log increase in IL-6: OR_adj_ 0.74, 95%CI: 0.68- 0.80; *P*= 0.086) (Supplementary Table 5).

### The causal effect of genetic liability to MDD on inflammatory markers

In reverse direction MR analyses, genetic liability to MDD was not associated with inflammatory markers (Standard deviation change [95% CI] per log odds higher liability to MDD: IL-6: - 0.001, 95%CI: −0.011 to 0.009; *P*=0.85; CRP: 0.003, 95%CI: −0.006 to 0.013; *P*=0.58; IL-1Ra: 0.003, 95%CI: −0.006 to 0.013; *P*=0.51). These findings were consistent in sensitivity analyses examining horizontal pleiotropy (Table 2).

**Table 2.** Causal estimates of genetic liability to major depressive disorder on circulating inflammatory markers levels

| Inflammatory marker | IVW<br>Beta (SE)<br>*P*-value | MR-Egger<br>Beta (SE)<br>*P*-value | Weighted median<br>Beta (SE)<br>*P*-value | Weighted mode<br>Beta (SE)<br>*P*-value |
| --- | --- | --- | --- | --- |
| CRP | 0.003 (0.005)<br>0.58 | -0.004 (0.14)<br>0.98 | 0.004 (0.006)<br>0.56 | 0.013 (0.015)<br>0.40 |
| IL1-Ra | 0.003 (0.005)<br>0.51 | -0.03 (0.14)<br>0.85 | 0.001 (0.006)<br>0.83 | -0.003 (0.013)<br>0.83 |
| IL-6 | -0.001 (0.005)<br>0.85 | -0.071 (0.140)<br>0.61 | -0.001 (0.006)<br>0.82 | -0.001 (0.014)<br>0.96 |

IVW = Inverse-variance weighted, SE= Standard error, CRP = C-reactive protein, IL1-Ra = Interleukin-1 Receptor antagonist, IL-6 = Interleukin-6. Beta represents the change in standard deviation units in levels of circulating inflammatory marker per natural log odds higher liability to major depressive disorder.

## DISCUSSION

In the present study we conducted two-sample MR to assess the bidirectional associations between circulating inflammatory markers and depression in 135,458 cases of major depressive disorder and 344,901 controls. We found evidence for a causal effect of interleukin-6 on risk of MDD. This effect was found to be independent of pleiotropic effects with neuroticism. In contrast to a previous MR analysis in UK Biobank^21^, we found little evidence for an association of C-reactive protein with risk of depression. Likewise, in contrast to some observational analyses, there was little evidence of association of interleukin-1 receptor antagonist with risk of MDD^10^. Finally, we found little evidence for an association of genetic liability to MDD on inflammatory markers.

### Evidence suggests IL-6 may influence risk of MDD

Our findings suggesting a causal role of circulating interleukin-6 in the development of major depressive disorder are consistent with insights from randomised trials along with findings from previous observational, genetic, and MR studies. A recent meta-analysis of 45 clinical studies investigating levels of serum inflammatory markers in depressed patients before and after treatment with currently licenced antidepressants found that antidepressants resulted in significant reductions in the levels of IL-6^41^. Additionally, *IL-6* genetic polymorphisms have been found to be associated with the outcomes of antidepressant treatment, and specifically relapse, remission time and treatment resistant depression^42^. Further, an emerging body of longitudinal studies have reported links between measured IL-6 levels and risk of depression across different populations^14,16,43,44^. Finally, a causal role of IL6 in depression is in agreement with a recent MR study investigating the associations between genetically proxied serum inflammatory markers and risk of depression in UK Biobank, suggesting a strong causal role of IL-6 in increasing the risk of probable lifetime major depression (moderate/severe) (OR per unit increase in log-transformed IL-6: 0.74, 95% CI: 0.62–0.89)^21^. It is important to note that there was some case overlap between our analysis and this previous UK Biobank analysis (there was approximately 10.5% case overlap in the UK Biobank)^21^. However, this degree of overlap is unlikely to have substantially contributed to the replication of this IL-6 finding. Furthermore, evidence from a previous MR study conducted in a birth cohort lends additional support for the importance of IL-6 in the aetiopathogenesis of depression^22^.

Free serum IL-6 is inactive as it cannot bind to gp130 unless bound to the IL-6 receptor. The lower circulating levels of IL-6 as proxied by genetic variants in *IL6R* used as instruments represents greater cellular binding of IL-6 and, thus, greater biological effects of IL-6^45^. Our findings implicating both circulating IL-6 and soluble IL-6 receptor thus suggest that the pharmacological targeting of IL-6 signalling through inhibition of the interleukin-6 receptor or interleukin-6 itself could be an effective therapeutic strategy for depression. Additionally, our findings suggest that intervening on upstream lifestyle or behavioural factors which influence depression risk, in part, through modulation of IL-6 levels could be an alternative risk-reducing strategy.

The exact mechanisms by which IL-6 may promote depression remains unclear, but several possibilities have been suggested. A meta-analysis of studies measuring serum cytokines in depression commented that all of the cytokines found to be elevated in depression (IL-1Ra, IL-6, TNFα and sIL2R) are modulated by the NFκβ pathway, which is one of the first pathways to respond to harmful cellular stimuli and activates many components of the anti-microbial response^46^. As described previously, IL-6 itself can activate two different signalling pathways, one of which appears to be anti-rather than pro-inflammatory^47^. The pro-inflammatory trans-signalling pathway is the principal pathway activated by IL-6 in the central nervous system^48^. Of note, evidence suggests that acute stress causes an inflammatory response which could explain why several disorders where stress is a risk factor are associated with the same immune alterations ^46,49^. Another mechanism to explain the increased risk with IL-6 is linked to the hypothalamus-pituitary-adrenal (HPA) axis. Glucocorticoid resistance, caused by the failure of the negative feedback loop in the HPA axis, is seen in 40-60% of individuals with depression^50^. The glucocorticoid receptor is known to inhibit NFκβ, which may increase the risk of an overactive inflammatory response in those with raised glucocorticoids levels such as individuals with depression^51^.

### The causal effect of CRP and IL-1Ra on risk of MDD

In the context of the present study, we found little evidence suggesting a causal effect of C-reactive protein on risk of major depressive disorder. This is in contrast to a recent MR analyses in UK Biobank suggesting a causal association between CRP and probable lifetime major depression (moderate/severe) (OR per unit increase in log-transformed CRP: 1.18, 95% CI: 1.07–1.29)^21^. However, an earlier Mendelian randomization analysis of 78,809 individuals from the Danish general population reported little evidence of an association of genetically proxied CRP on various depression-related phenotypes (e.g., hospitalisation with depression, prescription anti-depression medication use)^20^. Discrepancies among these findings could be attributed to differences in: i. sample size, ii. case definition and/or ascertainment, and iii. participant characteristics (e.g. age or severity of depression).

Furthermore, although IL-1Ra has been previously linked to depression in observational studies, we found limited evidence for a causal role in the development of MDD^46^. This could suggest confounding or chance but is unlikely to reflect reverse causation as the bidirectional analyses conducted in the present study provide evidence against a role of genetic liability to MDD on levels of IL-1Ra.

### The causal effect of MDD on inflammatory markers

In contrast to a number of observational studies suggesting bidirectional associations between inflammation markers and depression^15,16^, the present MR analysis found limited evidence to support a causal effect of genetic liability to MDD on inflammatory markers. These findings suggest that subclinical manifestations and/or pathophysiological mechanisms underpinning development of depression are unlikely to a large effect on levels of inflammatory markers examined.

### Strengths and Limitations

Strengths of this analysis include the use of a bidirectional Mendelian randomization approach which allowed us to: 1) clarify causal roles of inflammatory markers in depression, 2) estimate the magnitude of effect size, and 3) establish the likely direction of previously reported observational associations. Second, the construction of genetic instruments using either i) *cis*-acting variants or ii) multiple genome-wide significant variants allowed us to minimise horizontal pleiotropy and perform various sensitivity analyses to detect and/or adjust analyses for this bias, respectively. Third, the use of a two-sample MR approach allowed us to exploit summary genetic association data from several large genome-wide association studies, including the largest published MDD GWAS to date, which allowed us to substantially increase the number of depression cases included compared to the largest previous analysis (135,458 vs. 14,701 cases), in turn, increasing statistical power. Fourth, the restriction of inflammatory markers and MDD datasets to individuals of largely European ancestry reduced (but did not eliminate) confounding through population stratification in our analyses. The present findings are unlikely to be largely driven by residual confounding through population stratification.

There are several limitations to these analyses. First, controls within the Psychiatric Genetics Consortium were not all screened for mental health disorders, including depression. Whilst the potential inclusion of participants with depression as controls could bias inflammatory marker-depression analyses toward the null, advantages of using this approach include the ability to increase sample size which should mitigate such an attenuation. Second, although attempts were made to circumvent potential violations of MR assumptions in our analyses through the use of cis-acting variants as primary instruments and in sensitivity analyses, we cannot rule out the possibility of horizontal pleiotropy accounting for a causal role of IL-6 levels in development of depression. Likewise, we cannot rule out the possibility that false negative findings may have arisen through horizontally pleiotropic pathways biasing our findings toward the null. In addition, our analyses assumed linear associations between inflammatory markers and depression and assumed no interaction (e.g., gene-gene, gene-environment). Finally, the reverse direction analyses investigating the causal effects of genetic liability to MDD on circulating inflammatory markers had lower statistical power than analyses examining the role of inflammatory markers in depression.

## CONCLUSION

Within a two-sample MR framework we found evidence suggesting a causal role of IL-6 on risk of MDD in 135,458 cases and 344,901 controls in the Psychiatric Genetics Consortium. The identified causal effect was independent of pleiotropic effects with neuroticism. We were unable to replicate a previously reported association between genetically proxied CRP and depression. Further, there was little evidence to support a causal effect of CRP or IL-1Ra on MDD. In addition, there was little evidence to suggest bidirectional associations of reverse causation as an explanation for previously reported observational associations between CRP and IL-1Ra and depression. The present findings indicate the possible efficacy of interleukin-6 inhibition as a therapeutic target for depression and support the work currently ongoing in developing novel antidepressants targeting neuroinflammation, as well as trials repurposing already licensed anti-inflammatory agents e.g. celecoxib. Future research on the possible molecular and physiological pathways linking IL-6 and depression is expected to enhance current understanding on the underlying aetiopathogenic mechanisms of one of the leading causes of disability worldwide.

## Funding Acknowledgments

JY is part of the Medical Research Council Integrative Epidemiology Unit at the University of Bristol supported by the Medical Research Council [MC_UU_12013/1, MC_UU_12013/2, and MC_UU_12013/3]. CD is funded by the Wellcome Trust [108902/B/15/Z]. LS is funded by the David Telling Charitable Trust (E490). SJL is supported by the NIHR Biomedical Research Centre at the University Hospitals Bristol NHS Foundation Trust and the University of Bristol. No funding body has influenced data collection, analysis or its interpretation. This publication is the work of the authors, who serve as the guarantors for the contents of this paper. This work was carried out using the computational facilities of the Advanced Computing Research Centre - http://www.bris.ac.uk/acrc/ and the Research Data Storage Facility of the University of Bristol - http://www.bris.ac.uk/acrc/storage/.

## Conflict of interest

None

## References

1. de Graaf, R., Ten Have, M., van Gool, C. & van Dorsselaer, S. Prevalence of mental disorders and trends from 1996 to 2009. Results from the Netherlands Mental Health Survey and Incidence Study-2. Soc. Psychiatry Psychiatr. Epidemiol. 47, 203–213 (2012).

2. Takayanagi, Y. et al. Antidepressant use and lifetime history of mental disorders in a community sample: results from the Baltimore Epidemiologic Catchment Area Study. J. Clin. Psychiatry 76, 40 (2015).

3. Rush, A. J. et al. Acute and longer-term outcomes in depressed outpatients requiring one or several treatment steps: a STAR* D report. Am. J. Psychiatry 163, 1905–1917 (2006).

4. Machado-Vieira, R. et al. The timing of antidepressant effects: a comparison of diverse pharmacological and somatic treatments. Pharmaceuticals 3, 19–41 (2010).

5. Lex, H. et al. Quality of life across domains among individuals with treatment-resistant depression. J. Affect. Disord. 243, 401–407 (2019).

6. Kessler, R. C. The costs of depression. Psychiatr. Clin. 35, 1–14 (2012).

7. Smith, R. S. The macrophage theory of depression. Med. Hypotheses 35, 298–306 (1991).

8. Patten, S. B. Long-term medical conditions and major depression in a Canadian population study at waves 1 and 2. J. Affect. Disord. 63, 35–41 (2001).

9. Mikocka-Walus, A., Knowles, S. R., Keefer, L. & Graff L.Controversies revisited: a systematic review of the comorbidity of depression and anxiety with inflammatory bowel diseases. Inflamm. Bowel Dis. 22, 752–762 (2016).

10. Köhler, C. A. et al. Peripheral cytokine and chemokine alterations in depression: a meta□analysis of 82 studies. Acta Psychiatr. Scand. 135, 373–387 (2017).

11. Milaneschi, Y. et al. Interleukin-1 receptor antagonist and incident depressive symptoms over 6 years in older persons: the InCHIANTI study. Biol. Psychiatry 65, 973–978 (2009).

12. van den Biggelaar, A. H. J. et al. Inflammation and interleukin-1 signaling network contribute to depressive symptoms but not cognitive decline in old age. Exp. Gerontol. 42, 693–701 (2007).

13. Lassale, C. et al. Association of 10-Year C-reactive protein trajectories with markers of healthy aging: Findings from the English Longitudinal Study of Aging. Journals Gerontol. Ser. A 74, 195–203 (2018).

14. Khandaker, G. M., Pearson, R. M., Zammit, S., Lewis, G. & Jones, P. B. Association of serum interleukin 6 and C-reactive protein in childhood with depression and psychosis in young adult life: a population-based longitudinal study. JAMA psychiatry 71, 1121–1128 (2014).

15. Stewart, J. C., Rand, K. L., Muldoon, M. F. & Kamarck, T. W. A prospective evaluation of the directionality of the depression–inflammation relationship. Brain. Behav. Immun. 23, 936–944 (2009).

16. Huang, M. et al. Longitudinal association of inflammation with depressive symptoms: A 7-year cross-lagged twin difference study. Brain. Behav. Immun. 75, 200–207 (2019).

17. Davey Smith, G. & Ebrahim, S. ‘Mendelian randomization’: can genetic epidemiology contribute to understanding environmental determinants of disease? Int. J. Epidemiol. 32, 1–22 (2003).

18. Gage, S. H., Smith, G. D., Zammit, S., Hickman, M. & Munafò, M. R. Using Mendelian randomisation to infer causality in depression and anxiety research. Depress. Anxiety 30, 1185–1193 (2013).

19. Davies, N. M., Holmes, M. V & Smith, G. D. Reading Mendelian randomisation studies: a guide, glossary, and checklist for clinicians. Bmj 362, k601 (2018).

20. Wium-Andersen, M. K., Ørsted, D. D. & Nordestgaard, B. G. Elevated C-reactive protein, depression, somatic diseases, and all-cause mortality: a mendelian randomization study. Biol. Psychiatry 76, 249–257 (2014).

21. Khandaker, G. M. et al. Shared mechanisms between coronary heart disease and depression: findings from a large UK general population-based cohort. Mol. Psychiatry 1 (2019).

22. Khandaker, G. M., Zammit, S., Burgess, S., Lewis, G. & Jones, P. B. Association between a functional interleukin 6 receptor genetic variant and risk of depression and psychosis in a population-based birth cohort. Brain. Behav. Immun. 69, 264–272 (2018).

23. Davey Smith, G. & Hemani, G. Mendelian randomization: genetic anchors for causal inference in epidemiological studies. Hum. Mol. Genet. 23, R89–R98 (2014).

24. Wray, N. R. et al. Genome-wide association analyses identify 44 risk variants and refine the genetic architecture of major depression. Nat. Genet. 50, 668 (2018).

25. Haycock, P. C. et al. Best (but oft-forgotten) practices: the design, analysis, and interpretation of Mendelian randomization studies. Am. J. Clin. Nutr. 103, 965–978 (2016).

26. Pierce, B. L. & Burgess, S. Efficient design for Mendelian randomization studies: subsample and 2-sample instrumental variable estimators. Am. J. Epidemiol. 178, 1177–1184 (2013).

27. Ligthart, S. et al. Genome analyses of > 200,000 individuals identify 58 loci for chronic inflammation and highlight pathways that link inflammation and complex disorders. Am. J. Hum. Genet. 103, 691–706 (2018).

28. Matteini, A. M. et al. Novel gene variants predict serum levels of the cytokines IL-18 and IL-1ra in older adults. Cytokine 65, 10–16 (2014).

29. Sun, B. B. et al. Genomic atlas of the human plasma proteome. Nature. 558, 73 (2018).

30. Wald, A. The fitting of straight lines if both variables are subject to error. Ann. Math. Stat. 11, 284–300 (1940).

31. Burgess, S., Zuber, V., Valdes□Marquez, E., Sun, B. B. & Hopewell, J. C. Mendelian randomization with fine□mapped genetic data: Choosing from large numbers of correlated instrumental variables. Genet. Epidemiol. 41, 714–725 (2017).

32. Consortium, 1000 Genomes Project. A global reference for human genetic variation. Nature 526, 68 (2015).

33. Bowden, J., Davey Smith, G. & Burgess, S. Mendelian randomization with invalid instruments: effect estimation and bias detection through Egger regression. Int. J. Epidemiol. 44, 512–525 (2015).

34. Bowden, J., Davey Smith, G., Haycock, P. C. & Burgess, S. Consistent estimation in Mendelian randomization with some invalid instruments using a weighted median estimator. Genet. Epidemiol. 40, 304–314 (2016).

35. Hartwig, F. P., Davey Smith, G. & Bowden, J. Robust inference in summary data Mendelian randomization via the zero modal pleiotropy assumption. Int. J. Epidemiol. 46, 1985–1998 (2017).

36. Collaboration, I. G. C. E. R. F. Interleukin-6 receptor pathways in coronary heart disease: a collaborative meta-analysis of 82 studies. Lancet 379, 1205–1213 (2012).

37. Tyrrell, J. et al. Using genetics to understand the causal influence of higher BMI on depression. (2018).

38. Speed, D., Hemani, G., Speed, M. S., Børglum, A. D. & Østergaard, S. D. Investigating the causal relationship between neuroticism and depression via Mendelian Randomization. Acta Psychiatr. Scand. (2019).

39. Choi, K. W. et al. Assessment of bidirectional relationships between physical activity and depression among adults: a 2-sample mendelian randomization study. JAMA psychiatry (2019).

40. Sanderson, E., Davey, G. S., Windmeijer, F. & Bowden, J. An examination of multivariable Mendelian randomization in the single-sample and two-sample summary data settings. Int. J. Epidemiol. (2018).

41. Köhler, C. A. et al. Peripheral alterations in cytokine and chemokine levels after antidepressant drug treatment for major depressive disorder: systematic review and meta-analysis. Mol. Neurobiol. 55, 4195–4206 (2018).

42. Carvalho, S. et al. IL6-174G> C genetic polymorphism influences antidepressant treatment outcome. Nord. J. Psychiatry 71, 158–162 (2017).

43. Chu, A. L. et al. Longitudinal association between inflammatory markers and specific symptoms of depression in a prospective birth cohort. Brain. Behav. Immun. 76, 74–81 (2019).

44. Lamers, F. et al. Longitudinal Association Between Depression and Inflammatory Markers: Results From the Netherlands Study of Depression and Anxiety. Biol. Psychiatry (2019).

45. Scheller, J., Chalaris, A., Schmidt-Arras, D. & Rose-John, S. The pro-and anti-inflammatory properties of the cytokine interleukin-6. Biochim. Biophys. Acta (BBA)-Molecular Cell Res. 1813, 878–888 (2011).

46. Goldsmith, D. R., Rapaport, M. H. & Miller, B. J. A meta-analysis of blood cytokine network alterations in psychiatric patients: comparisons between schizophrenia, bipolar disorder and depression. Mol. Psychiatry 21, 1696 (2016).

47. Raison, C. L., Knight, J. M. & Pariante, C. Interleukin (IL)-6: a good kid hanging out with bad friends (and why sauna is good for health). Brain. Behav. Immun. (2018).

48. Hodes, G. E., Ménard, C. & Russo, S. J. Integrating Interleukin-6 into depression diagnosis and treatment. Neurobiol. Stress 4, 15–22 (2016).

49. Liu, Y.-Z., Wang, Y.-X. & Jiang, C.-L. Inflammation: the common pathway of stress-related diseases. Front. Hum. Neurosci. 11, 316 (2017).

50. Glassman, A. H. The dexamethasone suppression test: an overview of its current status in psychiatry. Am. J. Psychiatry 144, 1253–1262 (1987).

51. Pace, T. W. W., Hu, F. & Miller, A. H. Cytokine-effects on glucocorticoid receptor function: relevance to glucocorticoid resistance and the pathophysiology and treatment of major depression. Brain. Behav. Immun. 21, 9–19 (2007).

52. Dayawansa, N. H., Segan, J. D. S., Yao, H. H. I., Chong, H. I. & Sitzler, P. J. Incidence of normal white cell count and CJreactive protein in adults with acute appendicitis. ANZ J. Surg. 88, E539–E543 (2018).

53. Kleiner, G., Marcuzzi, A., Zanin, V., Monasta, L. & Zauli, G. Cytokine levels in the serum of healthy subjects. Mediators Inflamm. 2013, (2013).

