## Supplementary file for "Disentangling causal relationships between inflammatory markers and depression: a bidirectional Mendelian randomization analysis"

**SUPPLEMENTARY MATERIAL**

| **Contents** | | |
| --- | --- | --- |
| Page 2 | Supplementary Table 1 | Characteristics of genetic variants associated with inflammatory markers. |
| Page 4 | Supplementary Table 2 | Leave-one-out analyses for IL-6 instruments on risk of MDD. |
| Page 5 | Supplementary Table 3 | Causal estimates of IL-6 on risk of BMI, physical activity, neuroticism. |
| Page 6 | Supplementary Table 4 | Leave-one-out analyses for IL-6 instruments on neuroticism. |
| Page 7 | Supplementary Table 5 | The total and independent from neuroticism effects of IL-6 on risk of MDD (MVMR analysis). |
| Page 8 | Supplementary References | The GWAS studies utilized for the MR analyses investigating the effects of IL-6 on BMI, physical activity, neuroticism. |

**Supplementary Table 1. Characteristics of genetic variants associated with inflammatory markers**

| **Inflammatory marker** | **SNP** | **EA/NEA** | **EAF** | **Effect (SE)** | ***P*-value** |
| --- | --- | --- | --- | --- | --- |
| *CRP (mg/L)*  *(Liberal instrument)* |  |  |  |  |  |
|  | rs1051338 | G/T | 0.31 | 0.024 (0.004) | 9.87E-10 |
|  | rs10832027 | G/A | 0.33 | -0.026 (0.004) | 4.02E-11 |
|  | rs10838687 | G/T | 0.22 | -0.031 (0.004) | 4.59E-15 |
|  | rs112635299 | T/G | 0.02 | -0.107 (0.017) | 1.55E-10 |
|  | rs12202641 | T/C | 0.39 | -0.023 (0.004) | 4.46E-09 |
|  | rs12960928 | C/T | 0.27 | 0.024 (0.004) | 9.87E-10 |
|  | rs12995480 | T/C | 0.17 | -0.031 (0.005) | 2.82E-10 |
|  | rs1441169 | G/A | 0.53 | -0.025 (0.004) | 2.05E-10 |
|  | rs1490384 | T/C | 0.51 | -0.025 (0.004) | 2.05E-10 |
|  | rs1514895 | A/G | 0.71 | -0.027 (0.004) | 7.39E-12 |
|  | rs1558902 | A/T | 0.41 | 0.034 (0.004) | 9.48E-18 |
|  | rs1582763 | A/G | 0.37 | -0.022 (0.004) | 1.90E-08 |
|  | rs17658229 | C/T | 0.05 | 0.056 (0.01) | 1.07E-08 |
|  | rs178810 | T/C | 0.56 | 0.02 (0.004) | 2.87E-07 |
|  | rs1880241 | G/A | 0.48 | -0.028 (0.004) | 1.28E-12 |
|  | rs2239222 | G/A | 0.36 | 0.035 (0.004) | 1.07E-18 |
|  | rs2315008 | T/G | 0.31 | -0.023 (0.004) | 4.46E-09 |
|  | rs2352975 | C/T | 0.3 | 0.025 (0.004) | 2.05E-10 |
|  | rs2710804 | C/T | 0.37 | 0.021 (0.004) | 7.60E-08 |
|  | rs2836878 | G/A | 0.27 | 0.043 (0.004) | 2.96E-27 |
|  | rs2891677 | C/T | 0.46 | -0.02 (0.004) | 2.87E-07 |
|  | rs4092465 | A/G | 0.35 | -0.027 (0.004) | 7.39E-12 |
|  | rs4246598 | A/C | 0.46 | 0.022 (0.004) | 1.90E-08 |
|  | rs469772 | T/C | 0.19 | -0.031 (0.005) | 2.82E-10 |
|  | rs4774590 | A/G | 0.35 | -0.022 (0.004) | 1.90E-08 |
|  | rs6001193 | G/A | 0.35 | -0.028 (0.004) | 1.28E-12 |
|  | rs7121935 | A/G | 0.38 | -0.022 (0.004) | 1.90E-08 |
|  | rs75460349 | A/C | 0.97 | 0.086 (0.014) | 4.05E-10 |
|  | rs9271608 | G/A | 0.22 | 0.042 (0.005) | 2.23E-17 |
|  | rs9284725 | C/A | 0.24 | 0.027 (0.004) | 7.39E-12 |
| *CRP (mg/L)*  *(Conservative instrument)* |  |  |  |  |  |
|  | rs3093077 | C/A | 0.06 | 0.21 (0.0179) | 8.76E-32 |
|  | rs1205 | C/T | 0.67 | 0.18 (0.0102) | 1.07E-69 |
|  | rs1130864 | A/G | 0.30 | 0.13 (0.0077) | 5.99E-64 |
|  | rs1800947 | C/G | 0.94 | 0.26 (0.0153) | 9.18E-65 |
| *IL1Ra (pg/mL)* |  |  |  |  |  |
|  | rs6761276 | C/T | 0.58 | -0.1907 (0.0248) | 1.50E-14 |
| *IL6 (ng/mL)* |  |  |  |  |  |
|  | rs7529229 | T/C | 0.55 | 0.086 (0.012) | 6.15E-12 |
|  | rs4845371 | T/C | 0.43 | 0.062 (0.013) | 6.78E-07 |
|  | rs12740969 | T/G | 0.49 | 0.078 (0.013) | 2.2E-09 |
| *sIL6R* |  |  |  |  |  |
|  | rs2228145 | C/A | 0.39 | 0.2949 (0.015) | 1.8E-65 |

SNP = Single-nucleotide polymorphism, EA = Effect allele, NEA = Non-effect allele, EAF = Effect allele frequency, SE = Standard error

**Supplementary Table 2. Leave-one-out analyses for the IL-6 instruments on risk of MDD.**

| **SNP excluded** | **OR** | **95% CI** | ***P*-value** |
| --- | --- | --- | --- |
| rs7529229 | 0.86 | 0.72 – 1.02 | 0.086 |
| rs4845371 | 0.84 | 0.71 – 1.00 | 0.048 |
| rs12740969 | 0.83 | 0.69 - 1.00 | 0.050 |

SNP = Single-nucleotide polymorphism, OR = Odds ratio

**Supplementary Table 3. Causal estimates of IL-6 on risk of BMI, physical activity, neuroticism.**

|  | **Beta** | **95% CI** | ***P*-value** |
| --- | --- | --- | --- |
| Causal effect of IL-6 on BMI | -0.03 | -0.15 to 0.09 | 0.647 |
| Causal effect of IL-6 on physical activity | 0.46 | -0.31 to 1.23 | 0.240 |
|  | **OR** | **95% CI** | ***P*-value** |
| Causal effect of IL-6 on neuroticism | 1.07 | 1.02 to 1.12 | 0.011 |

OR = Odds ratio

**Supplementary Table 4. Leave-one-out analyses for the IL-6 instruments on risk of neuroticism.**

| **SNP excluded** | **OR** | **95% CI** | ***P*-value** |
| --- | --- | --- | --- |
| rs7529229 | 1.07 | 1.02- 1.13 | 0.007 |
| rs4845371 | 1.06 | 1.01- 1.12 | 0.019 |
| rs12740969 | 1.06 | 1.00- 1.12 | 0.039 |

OR = Odds ratio

**Supplementary Table 5. MVMR analysis investigating the causal effect of IL-6 on risk of MDD, adjusting for neuroticism.**

|  | **OR** | **95% CI** | ***P*-value** |
| --- | --- | --- | --- |
| Total Effects | 0.85 | 0.75- 0.96 | 0.007 |
| Independent Effects | 0.74 | 0.68- 0.80 | 0.086 |

OR = Odds ratio

**Supplementary References: The GWAS studies utilized for the MR analyses investigating the**

**effects of IL-6 on BMI, physical activity, neuroticism.**

Klimentidis YC, Raichlen DA, Bea J, Garcia DO, Wineinger NE, Mandarino LJ, et al. Genome-wide association study of habitual physical activity in over 377,000 UK Biobank participants identifies multiple variants including CADM2 and APOE. Int J Obes. 2018;42(6):1161–76.

Locke AE, Kahali B, Berndt SI, Justice AE, Pers TH, Day FR, et al. Genetic studies of body mass index yield new insights for obesity biology. Nature. Nature Publishing Group; 2015;518(7538):197.

Nagel M, Watanabe K, Stringer S, Posthuma D, van der Sluis S. Item-level analyses reveal genetic heterogeneity in neuroticism. Nat Commun. 2018;9(1):905.
